# Deficiency for scavenger receptors Stabilin-1 and Stabilin-2 leads to age-dependent renal and hepatic depositions of fasciclin domain proteins TGFBI and Periostin in mice

**DOI:** 10.1101/2023.02.09.526595

**Authors:** Thomas Leibing, Anna Riedel, Yannick Xi, Monica Adrian, Jessica Krzistetzko, Christof Kirkamm, Christof Dormann, Kai Schledzewski, Sergij Goerdt, Cyrill Géraud

## Abstract

**Background:** Stabilin-1 and Stabilin-2 are two major scavenger receptors of liver sinusoidal endothelial cells that mediate removal of diverse molecules from the plasma. Double-knockout mice (Stab-DKO) develop impaired kidney function and a decreased lifespan, while single Stabilin deficiency or therapeutic inhibition ameliorates atherosclerosis and Stab1-inhibition is subject of clinical trials in immuno-oncology. Although POSTN and TFGBI have recently been described as novel Stabilin ligands, the dynamics and functional implications of these ligands have not been comprehensively studied.

**Methods:** Immunofluorescence, Western Blotting and Simple Western™ as well as in situ hybridization (RNAScope™) und qRT-PCR were used to analyze transcription levels and tissue distribution of POSTN and TGFBI in Stab-KO mice. Stab-POSTN-Triple deficient mice were generated to assess kidney and liver fibrosis and function in young and aged mice.

**Results:** TGFBI and POSTN protein accumulated in kidney and liver tissue in Stab-DKO mice during aging despite unchanged transcriptional levels. Stab-POSTN-Triple KO mice showed glomerulofibrosis and a reduced lifespan comparable to Stab-DKO mice. However, alterations of the glomerular diameter and vascular density were partially normalized in Stab-POSTN-Triple KO.

**Conclusion:** TGFBI and POSTN are Stabilin-ligands that are deposited in an age-dependent manner in the kidneys and liver due to insufficient scavenging in the liver. Functionally, POSTN appears to partially contribute to the observed renal phenotype in Stab-DKO mice. This study provides details on downstream effects how how Stabilin dysfunction affects distant organs on a molecular and functional level.

## Introduction

During tissue turnover and regeneration, contents of the extracellular matrix, such as collagens and connective tissue polysaccharides are released into the plasma to a certain extent. Those molecules are cleared and degraded by scavenger receptors expressed by liver sinusoidal endothelial cells (LSECs) (Bhandari, Larsen, McCourt, Smedsrød, & Sørensen, 2021). Stabilin-1 (Stab1) and Stabilin-2 (Stab2) are the two members of the class H Scavenger Receptors expressed by LSECs, which contain EGF-like, laminin EGF-like (https://www.uniprot.org/uniprotkb/Q9NY15/entry) and fasciclin domains (PrabhuDas et al., 2017). They each have distinct, but also common ligands, based on the functionality of several domains (E. N. Harris & Cabral, 2019; Patten & Shetty, 2019). For example, hyaluronic acid is only bound by Stab2, while Stab1 is unable to bind hyaluronic acid, likely due to differences in link domain function (Hansen et al., 2005; Politz et al., 2002). Previously, we showed that Stab1/Stab2-double-knockout mice (Stab-DKO) present an impaired kidney function and a decreased lifespan likely due to their impaired scavenging function through LSECs (Schledzewski et al., 2011). Only recently, we were able to show that single Stabilin inhibition reduces atherosclerosis development and that extracellular matrix proteins Transforming growth factor, beta-induced, 68kDa (TGFBI) and Periostin (POSTN) are ligands of Stab1 and Stab2, likely due to fasciclin domain interactions and their plasma concentration is strongly increased in Stab-DKO mice (Manta et al., 2022). Interestingly, Stab1 and Stab2 as well as POSTN and TGFBI are the only four fasciclin-domain containing proteins in mammals (Seifert, 2018). POSTN is implicated as a causative agent in a plethora of fibrotic diseases: Knockout of POSTN in mice protects from CCl_4_-induced liver injury (Kumar et al., 2018) as well as unilateral ischemia-reperfusion injury induced kidney fibrosis (An et al., 2019). Additionally, it is implicated as a biomarker for disease severity in kidney disease (Guerrot et al., 2012; Satirapoj, Tassanasorn, Charoenpitakchai, & Supasyndh, 2015; Satirapoj et al., 2014; Wantanasiri, Satirapoj, Charoenpitakchai, & Aramwit, 2015). TGFBI is described as a modulator of cell-collagen interaction and thus seems likely to contribute to fibrous disease pathways. As TGFBI and POSTN constitute paralogs it seems probable that both proteins contribute in certain diseases (Schwanekamp, Lorts, Vagnozzi, Vanhoutte, & Molkentin, 2016). Since Stab1 and Stab2 single inhibition mediates beneficial effects in atherosclerosis development despite potentially inhibiting the scavenging of POSTN from the circulation and Stabilin double-deficiency damages kidney function, we here assess the role of POSTN in our models of genetic Stabilin deficiency.

## Methods

### Animals

All animal experiments were performed in accordance with local regulatory institutions (Regierungspräsidium Karlsruhe). All mice were bred on a C57BL/6J background in the animal facilities of the Centre for medical research, Medical Faculty Mannheim. Stab1-KO (B6.129S2-Stab1^tm1.1Cger^ (Schledzewski et al., 2011)) and Stab2-KO (B6.129S2-Stab2^tm1.1Cger^ (Schledzewski et al., 2011)) mice were interbred to generate transgenic mice with Stabilin-double deficiency (Stab-DKO). Stab-DKO mice were bred to POSTN-KO mice (B6;129-Postn^tm1Jmol^/J (Oka et al., 2007)) to generate Stab-POSTN-Triple deficient mice (Stab-POSTN-TrKO). Male and female mice were pooled for analysis except in **Supp. Fig. 1.**

### Immunofluorecence

PFA-fixed and paraffinized liver and kidney tissue was sectioned at 4 microns. Deparaffinization, rehydration and antigen retrieval were performed as previously described (Leibing et al., 2018). Primary antibodies **(Supp. Table 1)** were incubated overnight and secondary antibodies (DIANOVA) for one hour.

### Simple Western

Simple Western was performed according to manufacturer’s protocols (V. M. Harris, 2015; Lück, Haitjema, & Heger, 2021)). Primary antibodies are found in **Supp. Table 1**.

### Western blotting

Kidney and liver tissue were homogenized in RIPA and were analysed by SDS-Page and Immunoblotting on PVDF membranes as by the manufacturer protocol (Trans-Blot Turbo Transfer System, Bio-Rad). Incubation with primary antibodies **(Supp. Table 1)** was performed at 4°C overnight at a 1:1000 dilution in 5% nonfat dry milk (Bio-Rad No. 1706404). Incubation of HRP conjugated secondary antibody (different species, Dianova) was performed for 1 hour at room temperature. Chemiluminescence (Millipore WBLUF0500) intensity detection was performed with the Azure^®^ c400 imaging system (Azure ^®^ Biosystems) and quantified using ImageJ 1.53 (Schneider, Rasband, & Eliceiri, 2012).

### RNA in situ hybridization

PFA-fixed and paraffinized liver and kidney tissue was sectioned at 4 μm. A modified non-isotopic in-situ hybridization protocol was carried out using the RNAscope 2.5 HD Red kit (Advanced Cell Diagnostics) following the manufacturer’s recommended protocol with specific probes against the positive control mouse Ppib (Cyclophilin B) gene and mouse POSTN or TGFBI **(Supp. Table 1)**. Sections were ultimately stained with DAB and counterstained with hematoxylin.

### Urinalysis

Animals of the same genotype were arranged in groups of 6 to 8 and followed for an observation period of up to 2 years. Mice were housed in metabolic cages for 24 hours to collect urine samples. Urine was analyzed for Protein and Albumin (Cobas c 311 Analyzer; Roche Diagnostics).

### Statistics

Statistical analyses were performed with JMP^®^ 16 and SAS^®^ 9.4M7 (SAS Institute Inc.). For comparisons between two groups, t-test was used when the normality assumption was met using the Shapiro-Wilk test. For group comparisons of three groups, the one-way ANOVA was used when the normality assumption was met using the Shapiro-Wilk test. We used the Brown-Forsythe test to check for equal variances. In case of unequal variances, the Welch ANOVA was used. Tukey-Kramer HSD was used as a post-hoc tests for groups of three. For group comparisons of four or more groups, the one-way ANOVA was used when the normality assumption was met using the Shapiro-Wilk test. We used the Brown-Forsythe test to check for equal variances. In case of unequal variances, the Welch ANOVA was used. For post-hoc analyses after significant ANOVA p-values, Dunnett’s test was used. If the normality assumption was not met, we used a Kruskal-Wallis test followed by Steel’s multiple comparison test. Reference groups for Dunnett’s test or Steel with control were WT mice. Differences between data sets with P < 0.05 in t-test, ANOVA, Kruskal-Wallis test and post-hoc tests were considered statistically significant. For comparison of old and young animals, Standard Least Squares with emphasis on Effect leverage was used. Effects of age, genotype and a full factorial analysis of Age*Genotype was performed. Test slices were used to compare different ages in a single genotype.

### Image quantification

Signal intensities were quantified in labeled glomerular area or total liver area using thresholding in ImageJ 1.53 (Schneider et al., 2012).

### Rt-PCR

Reverse transcription (RT) into complementary DNA (cDNA) was conducted using Maxima Reverse Transcriptase (EP0752, Thermo Fisher Scientific) and Oligo(dT)18 primer (SO131, Thermo Fisher Scientific). Quantitative PCR (qPCR) was performed using innuMIX qPCR SyGreen Sensitive (845-AS-1310200, Analytik Jena, Jena, Germany) on a qTOWER 3 G touch thermal cycler (Analytik Jena). Primers for RT-qPCR were designed using NCBIs PrimerBLAST (https://www.ncbi.nlm.nih.gov/tools/primer-blast/). If possible, primer pairs **(Supp. Table 1)** were designed to span an exon-exon junction, thus being mRNA-specific. If this was not possible they needed to be separated by at least one intron. Primers were validated using no template controls, no RT controls, agarose gel electrophoresis and melt curve analysis. Amplification data were analyzed using qPCRsoft 4.0.8.0 (Analytik Jena). Normalized expression values were calculated using the Pfaffl method considering amplification efficiency values determined by standard curves.

## Results

### Stabilin ligand POSTN is more abundant in aged, enlarged Stab-DKO glomeruli and liver tissue compared to WT

Immunofluorescence revealed an increased abundance of Stabilin-ligand POSTN in glomeruli from aged (12 months old) Stab-DKO mice compared to aged WT mice **(Fig. 1A)**. Quantification revealed Stab1-KO and Stab2-KO glomeruli did not show significant alterations in POSTN abundance compared to WT **(Fig. 1B)**. Stab-DKO glomeruli were significantly larger than WT glomeruli, Stab1-KO and Stab2-KO glomeruli diameter did not reach significance **(Fig. 1C)**. Furthermore, perisinusoidal POSTN staining in Stab-DKO livers was observed, which was absent in WT mice **(Fig. 1D)**.

**Fig. 1.**
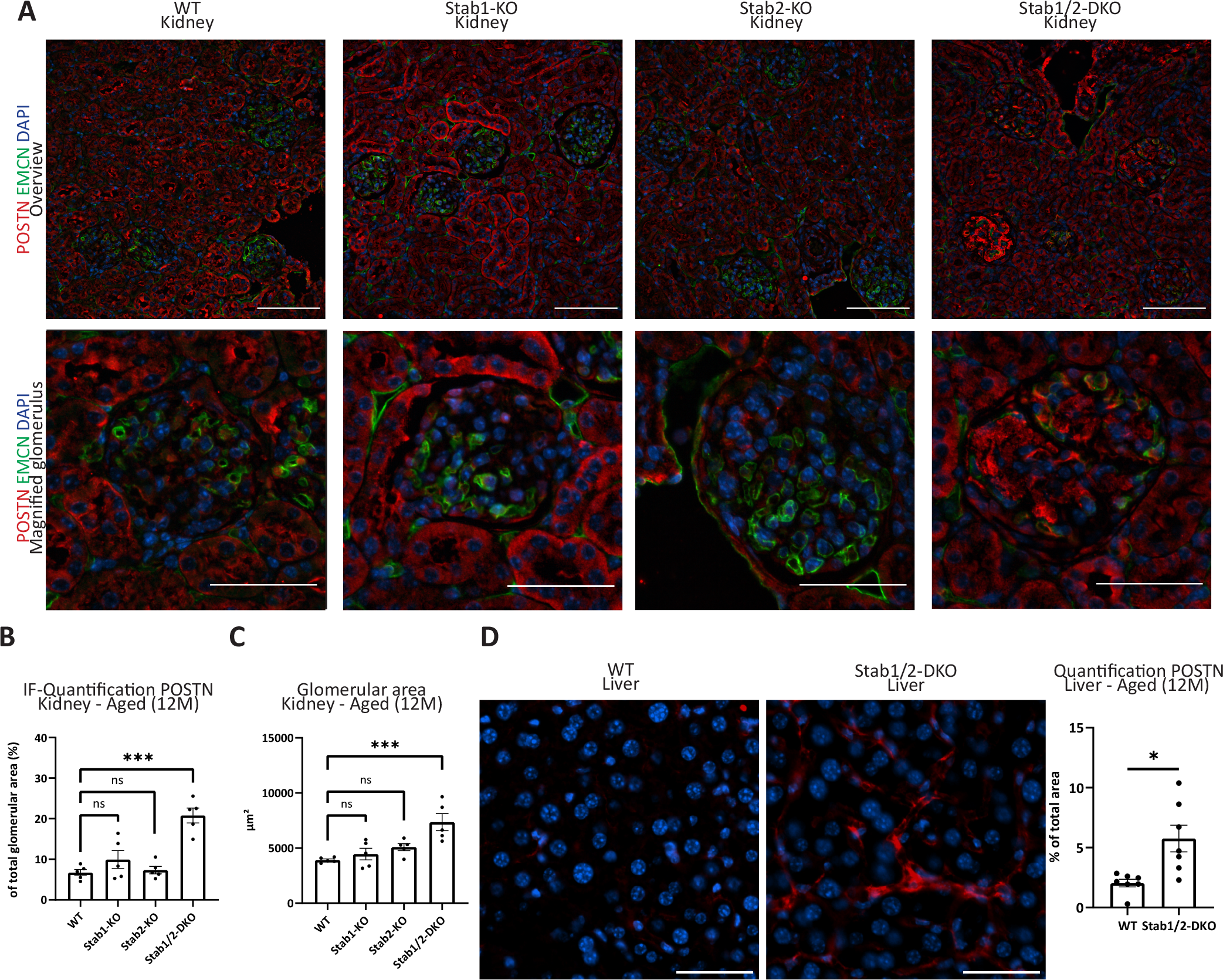
POSTN in kidneys and liver from Stabilin-deficient animals. **(A)** Representative photomicrographs of kidney tissue (upper lane: Overview; lower lane: Magnified glomeruli) co-stained with DAPI (blue), EMCN (green) and POSTN (red). **(B)** Quantification of average POSTN-positive area in glomeruli (in % of total glomerular area). **(C)** Quantification of average glomerular diameter (in μm^2^). **(D)** Representative photomicrographs of liver tissue co-stained with DAPI (blue) and POSTN (red). Quantification of average POSTN-positive area of total photomicrograph area is shown on the right (in % of total photomicrograph area). Scale bar = 50 μm. n≥5 for all experiments. ns = not significant, *= p<0.05; **=p<0.01; ***=p<0.001.

### POSTN expression is unaltered in Stab-DKO tissues, indicating deposition from plasma

To explore the mechanism of increased POSTN abundance, we performed RNA-ISH for POSTN in glomeruli and liver tissue of aged Stab-DKO mice in comparison to WT mice. Furthermore, we performed qRT-PCR of kidney lysate and liver lysate. Here, no difference in the RNA staining pattern and expression levels in kidney and liver **(Fig. 2)** were found, indicating no increased transcription and thereby no increased production of POSTN in both organs in Stab-DKO mice, suggesting a deposition due to increased plasma levels.

**Fig. 2.**
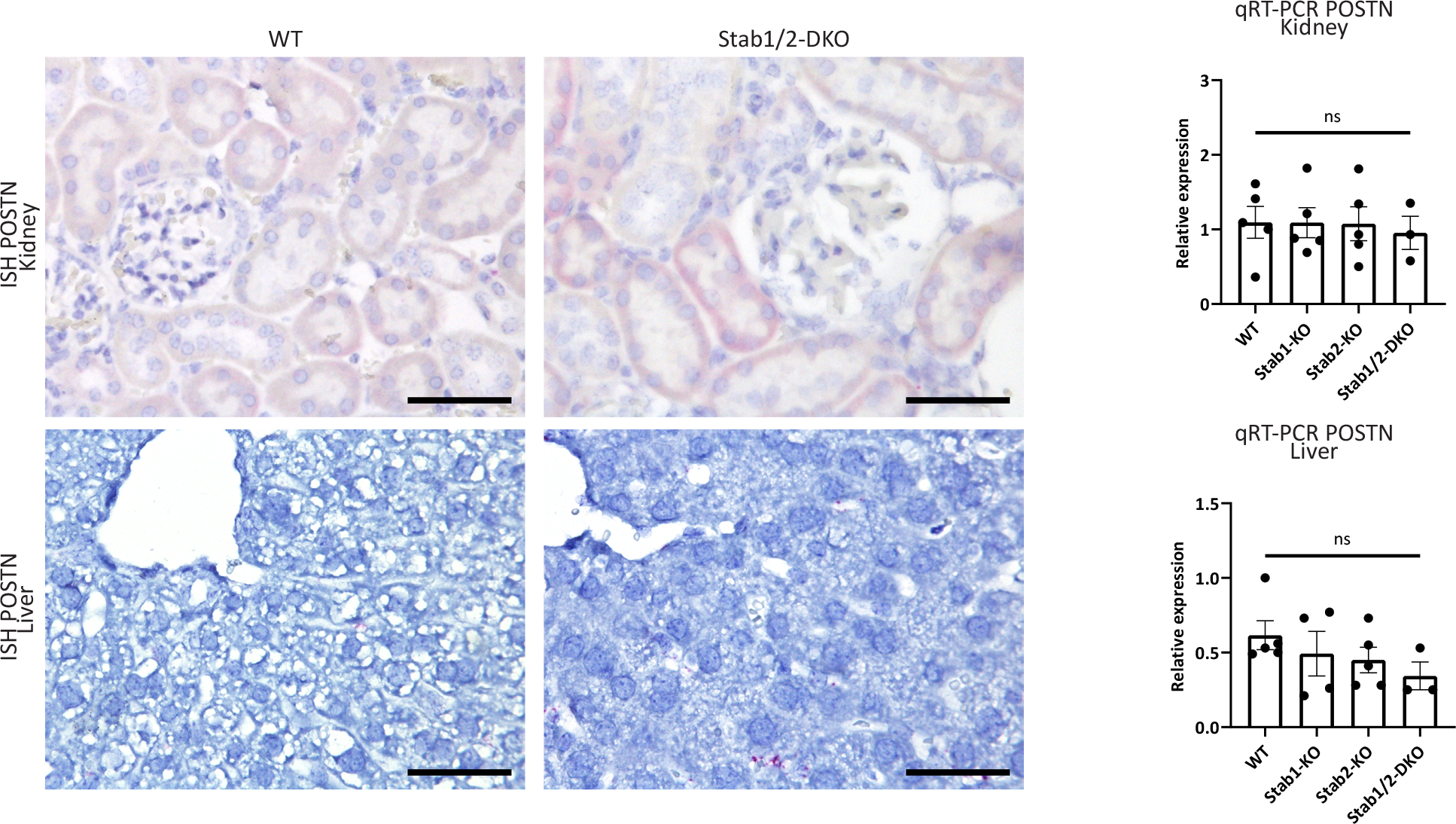
Local POSTN transcription is unaltered despite increased protein levels. Left panel: In-situ hybridization and Right graph: rt-PCR of Kidney (upper panels) and Liver (lower panels) in WT and Stab-DKO animals for POSTN. Scale bar = 50 μm. n≥3 for all experiments. ns = not significant, *= p<0.05; **=p<0.01; ***=p<0.001.

### Genetic ablation of POSTN does not rescue shortened lifespan, glomerulofibrosis and albuminuria observed in Stab-DKO mice

To assess whether POSTN depositions in Stab-DKO glomeruli and liver tissue might be causative for the observed glomerulofibrotic nephropathy and reduced lifespan in Stab-DKO mice, we crossed POSTN-KO mice with Stab-DKO mice. Stab-POSTN-TrKO mice were viable but showed a similarly shortened lifespan comparable to Stabilin-DKO mice **(Fig. 3A)**. Age-related mortality was comparable between male and female mice **(Suppl. Fig. 1)**. Urinalysis of aged WT, Stab-DKO and Stab-POSTN-TrKO mice revealed comparable proteinuria and albuminuria in Stab-DKO and Stab-POSTN-TrKO which was absent in WT mice **(Fig. 3B)**. POSTN staining was not observed in Stab-POSTN-TrKO glomeruli and livers serving as negative control supporting the specificity of our previous findings **(Supp. Fig. 2A)**.

**Fig. 3.**
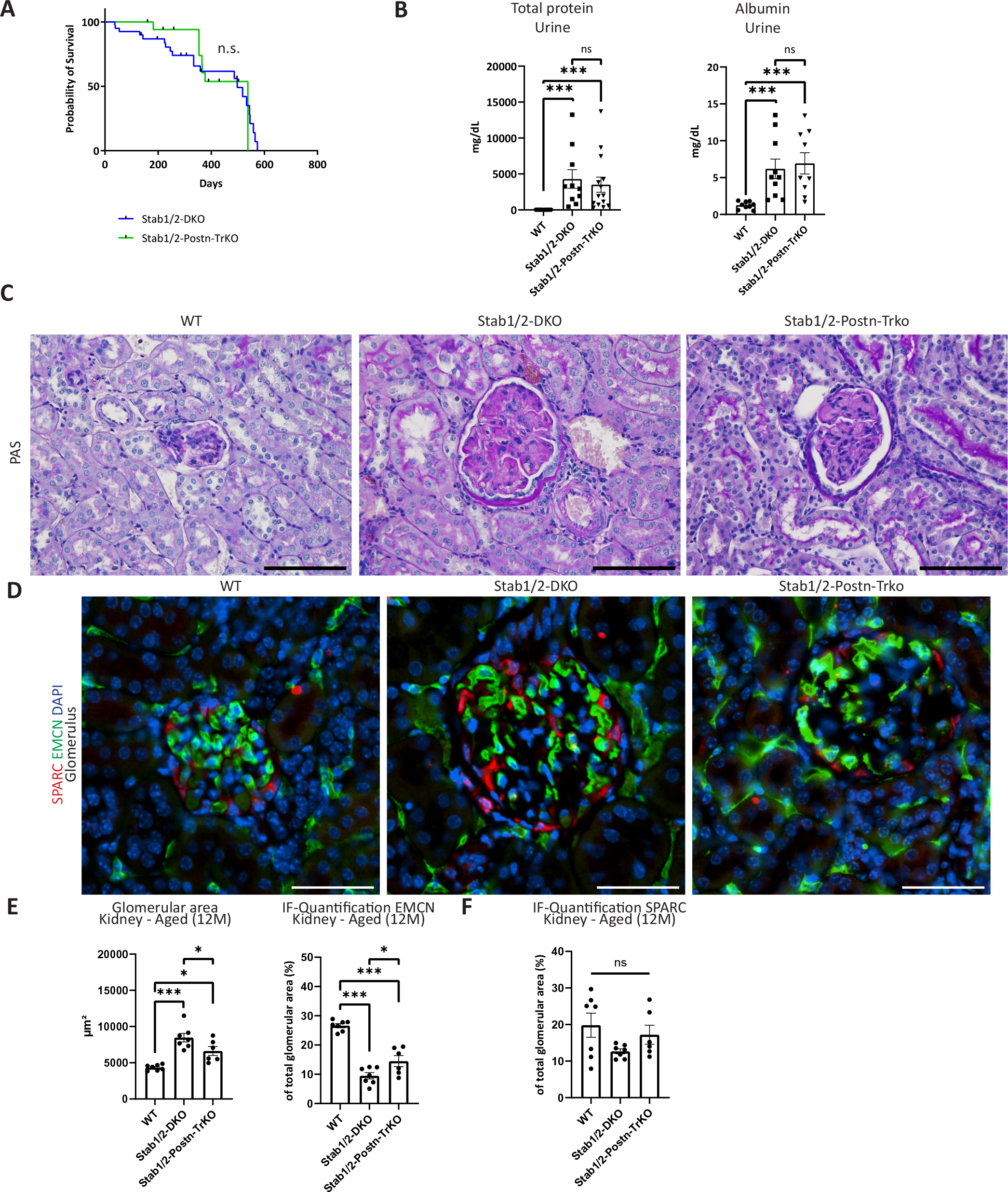
Ablation of POSTN does not influence phenotype in Stab-DKO mice. **(A)** Survival curve of Stab-DKO mice and Stab-POSTN-Triple deficient mice. **(B)** Total protein and Albumin levels in different mice strains were assessed **(C)** Representative photomicrographs of PAS-stained kidney tissue from different mice strains. **(D)** Representative photomicrographs of kidney tissue co-stained with DAPI (blue), EMCN (green) and SPARC (red). **(E)** Quantification of glomerular diameter and EMCN-positivity in total glomerular area, **(F)** Quantification of SPARC-positivity in total glomerular area. Scale bar = 50 μm. n≥5 for all experiments. ns = not significant, *= p<0.05; **=p<0.01; ***=p<0.001.

To further explore possible phenotypic alterations in Stab-POSTN-Triple deficient mice, PAS staining of kidneys from aged WT, Stab-DKO and Stab-POSTN-TrKO were performed. No obvious differences between PAS-positivity in glomeruli Stab-DKO and Stab-POSTN-TrKO wwas observed, WT glomeruli did not display large amounts of PAS-positive material. Stab-POSTN-TrKO appeared slightly smaller than Stab-DKO glomeruli **(Fig. 3C)**. To conform our findings, we quantified the glomerular area, vascular density (using EMCN as a surrogate) and podocyte coverage (using SPARC as a surrogate) in immunofluorescence stainings for DAPI, EMCN and SPARC **(Fig. 3D)** which revealed enlarged glomeruli with a reduced vascularized area in Stab-DKO and Stab-POSTN-TrKO compared to WT **(Fig. 3E)**. Slight differences were found between Stab-DKO and Stab-POSTN-TrKO, as Stab-POSTN-TrKO showed a smaller glomerular are and a higher vascularized glomerular area compared to Stab-DKO **(Fig. 3E)**. Interestingly, Podocyte marker SPARC showed a similar staining pattern and abundance relative to glomerular area in all genotypes, indicating a lack of podocyte involvement in the type of glomerulofibrosis and reduced vascularization observed in Stab-DKO and Stab-POSTN-TrKO mice **(Fig. 3E-F)**.

Liver fibrosis levels were scored using Sirius-red quantification, which did not show significant differences in Sirius-red positive areas between Stab-DKO and Stab-POSTN-TrKO mice but were elevated compared to WT in both knockout mice lines **(Supp. Fig. 2B)**.

### Stabilin ligand TGFBI is highly abundant in diseased glomeruli and livers from Stabilin-deficient animals

Since genetic deletion of POSTN did not rescue the phenotype of double-Stabilin deficiency, we assessed abundance of another newly identified Stabilin-ligand containing fasciclin domains, TGFBI, in glomeruli of Stabilin-deficient animals. Interestingly, TGFBI showed a similar pattern in immunofluorescent stainings in Stab-DKO similar to POSTN both in kidneys **(Fig. 4A**, **Supp. Fig. 3)** and liver **(Supp. Fig. 3)**. Quantification of TGFBI intensity in aged glomeruli revealed significantly more TGFBI in Stab-DKO glomeruli compared to WT, while Stab1-KO and Stab2-KO glomeruli did not exhibit significantly more TGFBI **(Fig. 4B)**. Similarly to POSTN, we performed RNA-ISH for TGFBI in glomeruli of aged Stab-DKO mice in comparison to WT mice as well as qRT-PCR of kidney lysate which did not show any differences, pointing towards a deposition of TGFBI from the plasma **(Fig. 4C)**.

**Fig. 4.**
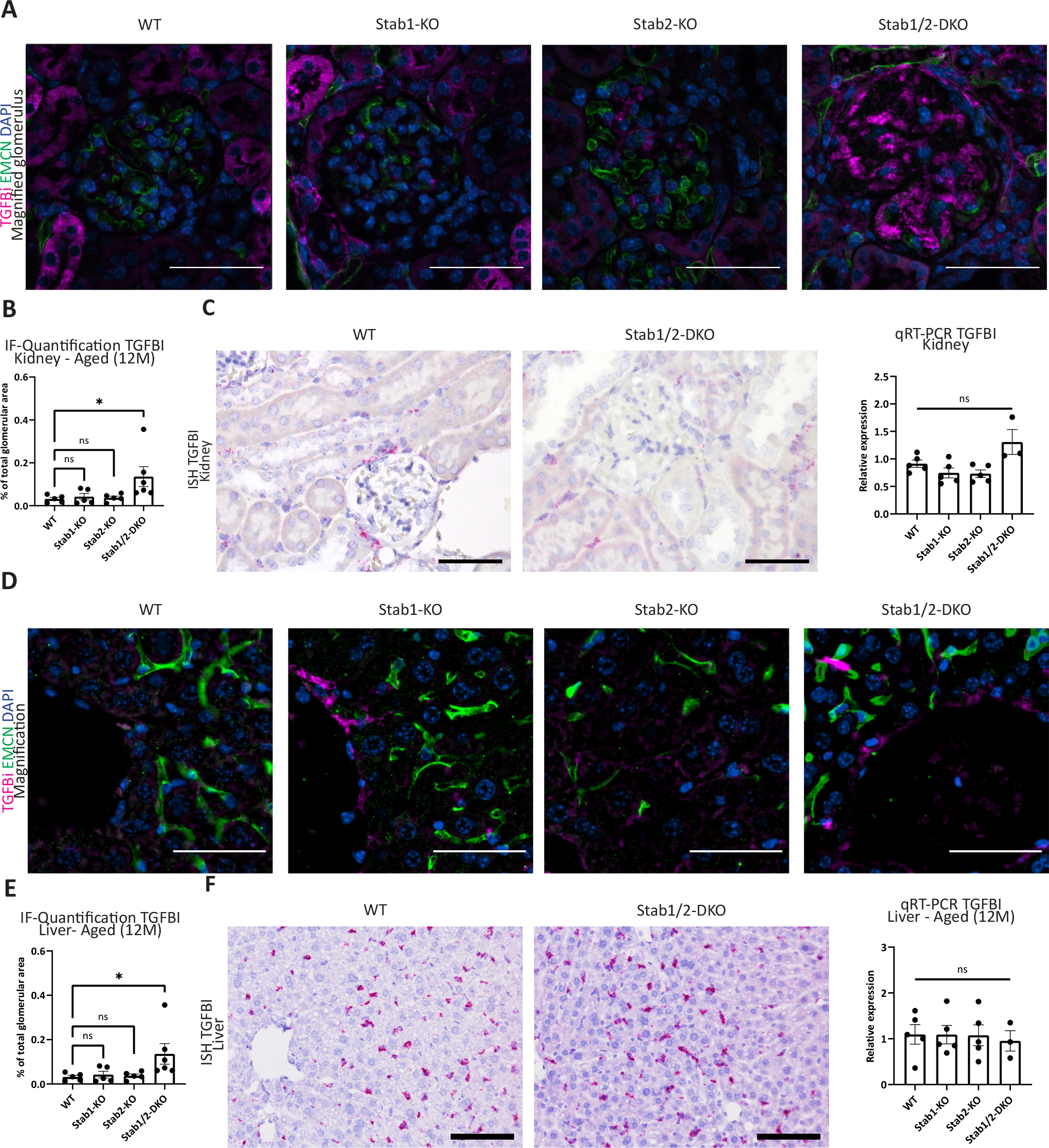
Abundance of TGFBI in organs from Stabilin-deficient animals. **(A)** Representative photomicrographs of kidney tissue (upper lane: Overview; lower lane: Magnified glomeruli, Scale bar = 50 μm) co-stained with DAPI (blue), Emcn (green) and TGFBI (magenta). **(B)** Quantification of average TGFBI-positive area in glomeruli (in % of total glomerular area, n≥5). **(C)** Left panel: In-situ hybridization (Scale bar = 50 μm) and right graph: rt-PCR of Kidney tissue in WT and Stab-DKO animals for TGFBI (n≥3). **(D)** Representative photomicrographs of liver tissue (upper lane: Overview; lower lane: Magnified pericentral area, Scale bar = 50 μm) co-stained with DAPI (blue), Emcn (green) and TGFBI (magenta). **(E)** Quantification of average TGFBI-positive area of total photomicrograph area (in % of total photomicrograph area, n≥5). **(F)** Left panel: In-situ hybridization (Scale bar = 100 μm) and Right graph: rt-PCR (n≥3) of Kidney tissue in WT and Stab-DKO animals for TGFBI. ns = not significant, *= p<0.05; **=p<0.01; ***=p<0.001.

Since POSTN was found to be more abundant in Stab-DKO livers, we also analyzed liver tissue for TGFBI abundance. Here, similarly to the kidney, we found a pronounced increase in TGFBI levels only in Stab-DKO livers compared to WT in immunofluorescence **(Fig. 4D-E)** and no increased transcription using RNA-ISH and qRT-PCR **(Fig. 4F)**.

To confirm antibody specificity, we performed Simple-Western **(Supp. Fig. 4A)** and Western Blotting **(Supp. Fig. 4B)**, which revealed increased TGFBI abundance in kidney and liver lysate of Stab-DKO animals. No overt changes were observed in Stab1-KO und Stab2-KO liver and kidney tissue in comparison to WT mice, although a small but significant increase was found in Stab2-KO livers using simple Western, which could not be confirmed with other methods.

### TGFBI depositions occur independently of POSTN in Stab-DKO tissue

Previously, we have demonstrated that POSTN and TGFBI are both ligands of Stab1 and Stab2, likely due to their structure and binding of fasciclin domains. Since TGFBI and POSTN revealed similar staining patterns in Stab-DKO animals, we correlated TGFBI and POSTN abundance in single glomeruli and found a strong and significant correlation in Stab-DKO, but not WT glomeruli **(Fig. 5A)**. As TGFBI and POSTN may directly interact via there Fasciclin domains, we checked whether increased TGFBI abundance in glomeruli and liver tissue occurs independently of POSTN in Stabilin-DKO mice. Comparison of TGFBI abundance in WT, Stab-DKO and Stab-POSTN-TrKO kidney and liver tissue revealed increased TGFBI abundance in Stab-DKO and Stab-POSTN-TrKO compared to wildtype control mice, indicating independence of TGFBI abundance in kidney **(Fig. 5B)** and liver tissue **(Fig. 5C)** in Stab-DKO mice from POSTN.

**Fig. 5.**
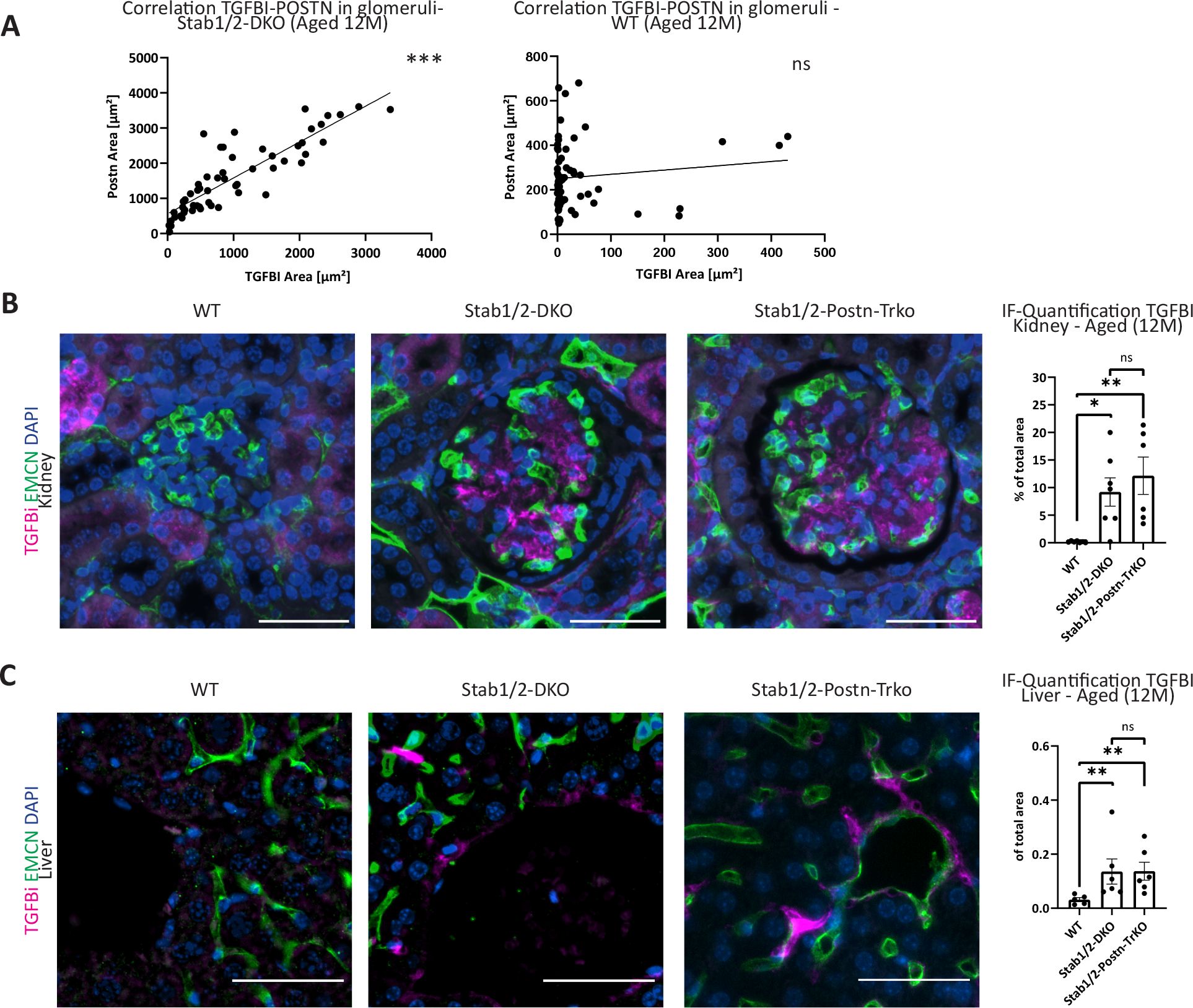
Stab-TrKO mice share similar TGFBI depositions in liver and kidney with Stab-DKO mice. **(A)** Correlation of TGFBI and POSTN abundance from quantified immunofluorescent microphotographs in kidney tissue of DKO (left panels) and WT (right panels) mice. **(B)** Representative photomicrographs of kidney tissue co-stained with DAPI (blue), Emcn (green) and TGFBI (magenta). Right panel: Quantification of average TGFBI-positive area in glomeruli (in % of total glomerular area). **(C)** Representative photomicrographs of liver tissue co-stained with DAPI (blue), Emcn (green) and TGFBI (magenta). Right panel: Quantification of average TGFBI-positive area of total photomicrograph area (in % of total photomicrograph area). n≥5 for all experiments. Scale bar = 50 μm. ns = not significant, *= p<0.05; **=p<0.01; ***=p<0.001.

### POSTN and TGFBI positivity in liver and kidney is detected in early adult age in Stab-DKO and increases during aging

Since we could show strong positivity of TGFBI and POSTN in aged Stab-DKO mice which already show a pronounced proteinuria, we investigated whether 3-month-old animals show similar staining patterns in the kidney and liver. Here, we observed a strong increase of POSTN **(Fig. 6A)** and TGFBI **(Fig. 6B)** positive area in glomeruli of young Stab-DKO mice compared to young WT controls. Age-matched Stab1-KO and Stab2-KO glomeruli did not show any differences in POSTN-positive area, a slight increase in TGFBI-positive area was observed in Stab2-KO glomeruli. Comparing glomeruli from young mice to aged mice revealed that ligand abundance did not change in WT and Stab2-KO mice, while Stab1-KO mice showed significantly more POSTN positive area in aged glomeruli and Stab-DKO mice showed an age-dependent increase for both TGFBI and POSTN.

**Fig. 6.**
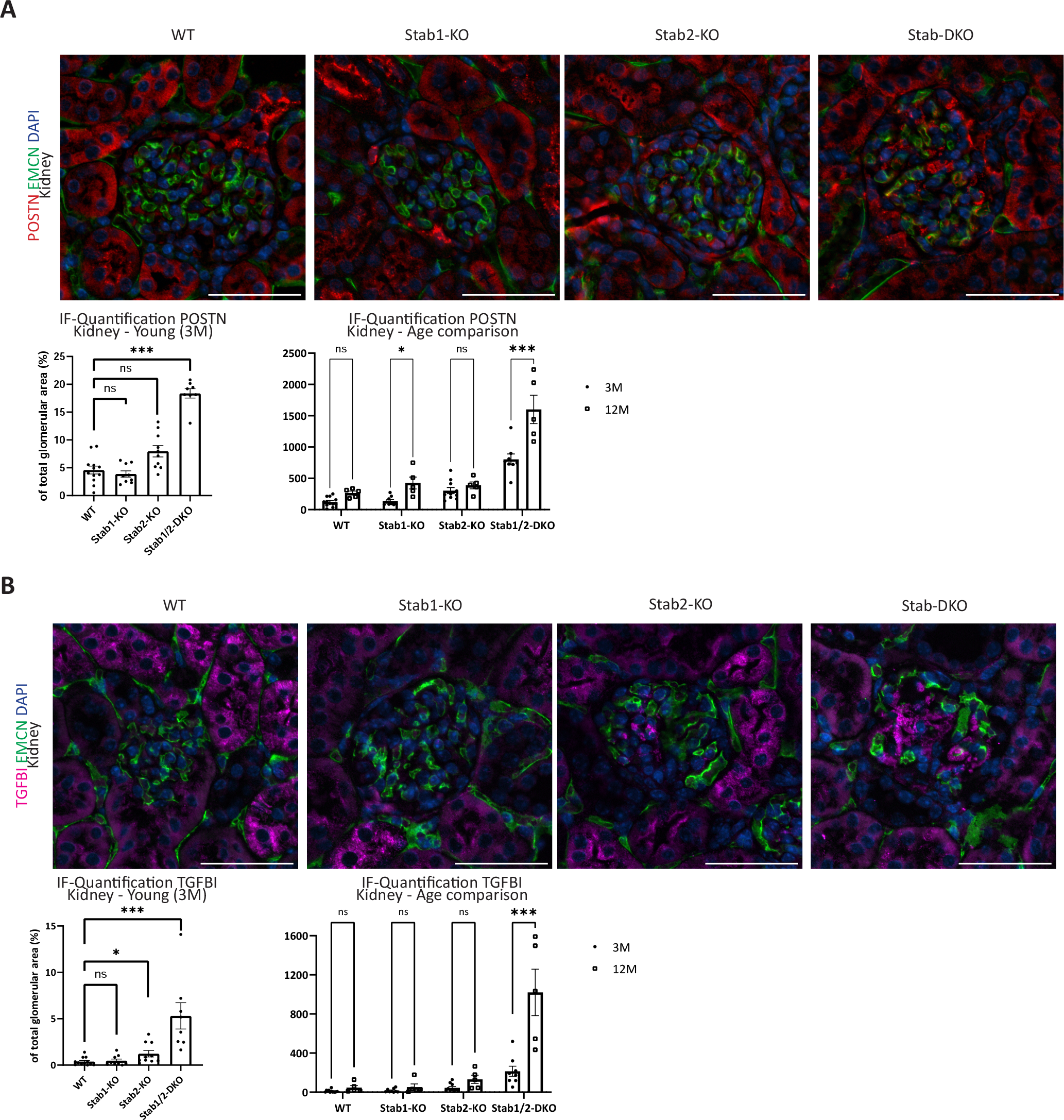
Age-dependent ligand abundance in glomeruli. **(A)** Representative photomicrographs of kidney tissue co-stained with DAPI (blue), Emcn (green) and POSTN (red). Lower left panel: Quantification of average POSTN-positive area in glomeruli (in % of total glomerular area). Lower right panel: Glomerular POSTN abundance from quantified immunofluorescent microphotographs in 3- and 12-months old mice. **(B)** Representative photomicrographs of kidney tissue co-stained with DAPI (blue), Emcn (green) and TGFBI (magenta). Lower left panel: Quantification of average TGFBI-positive area in glomeruli (in % of total glomerular area). Lower right panel: Glomerular TGFBI abundance from quantified immunofluorescent microphotographs in 3- and 12-months old mice. n≥5 for all experiments. Scale bar = 50 μm. ns = not significant, *= p<0.05; **=p<0.01; ***=p<0.001.

## Discussion

We here show that scavenging by Stab1 and Stab2 is crucial not only for homeostasis of TGFBI and POSTN plasma levels in mice (Manta et al., 2022), but also to prevent age-dependent accumulation of TGFBI and POSTN in distant organs. We hypothesize that POSTN and TGFBI are not produced locally in Stab-DKO livers and kidneys, but are deposited in distant organs, possibly due to their high concentrations in Stab-DKO plasma, since their scavenger receptors are missing. This hypothesis is supported by constant mRNA levels in the liver and kidney as shown here by RNA-ISH and qRT-PCR. As we have shown previously, transcriptomic data from isolated LSECs show a slight but significant downregulation of POSTN in LSECs from Stabilin deficient animals compared to WT animals (Olsavszky et al., 2021). POSTN and TGFBI staining was confined to the perisinusoidal area in the liver, with a similar pattern compared to Sirius-red positive areas, indicating a colocalization with collagens in the Space of Disse.

In mice and humans, there are somewhat contradictory findings regarding age-dependent plasma levels of TGFBI and POSTN. In mice, it was described that TGFBI and POSTN plasma levels are decreased in aged mice (Yang et al., 2020), while human data show an increase in aged populations (Lehallier et al., 2019). Since aging in mice and humans differ considerably, we can only speculate on the biological relevance of decreased TGFBI and POSTN plasma abundance in aged WT mice. Since we were able to find increased deposition of both Stabilin ligands in livers and kidneys of aged knockout mice compared to younger mice, we could further underline the hypothesis of a time-dependent deposition from plasma in our models.

Interestingly, ablation of POSTN in Stab-DKO mice alone does not seem to fully rescue the reduced lifespan or kidney defects observed in these mice, although POSTN is heavily implicated mechanistical development of liver and kidney fibrosis as described earlier. Knockout of POSTN also did not affect levels of TGFBI in glomeruli and livers of Stab-DKO mice, indicating that TGFBI depositions occur independently of POSTN. In comparison to Stab-DKO, in average glomerular diameter and vascularization in Stab-POSTN-TrKO were observed, which did not influence survival or protein levels in urine. Nevertheless, these findings point towards a slight amelioration of the kidney phenotype by deletion of POSTN. Whether TGFBI is solely responsible for the possibly fatal glomerulofibrosis in Stab-DKO and Stab-POSTN-TrKO is unclear, since we observed more than 100 dysregulated proteins comparing the plasma proteome of Stab-DKO mice compared to WT mice (Manta et al., 2022). Stab1 and Stab2 not only contain fasciclin domains, but also EGF-like domains and Laminin EGF-like domains beside the X-link domain (Politz et al., 2002). The EGF-like domain containing protein Reelin has previously been identified by us as a Stabilin-2 ligand (Manta et al., 2022). No Reelin depositions were found in kidneys and livers of our models (data not shown). Other proteins elevated in the plasma of Stabilin deficient mice containing EGF-like domains and Laminin EGF-like domains are putative ligands of Stabilin receptors and will be scrutinized in further experiments.

These findings bear consequences for the use of therapeutical inhibition of Stabilins, which for Stab1 is already in Phase 2 clinical trials in cancer (Virtakoivu et al., 2021) and might be a treatment option for atherosclerosis in future studies (Manta et al., 2022). Single deficiency for Stab1 or Stab2 alone did not seem to induce strong damage to kidney function, although a mild increase in TGFBI depositions was observed in young, but not aged Stab2-KO glomeruli. It can be assumed that absence of one of the Stabilins can be compensated for by the respective other Stabilin and only double-knockout is detrimental, which is in line with previous findings from our group (Manta et al., 2022; Schledzewski et al., 2011). In summary, we do not expect grave side effects form antibody-mediated Stabilin-inhibition, which is usually transient and not as strong as genetic deficiency for one or even both Stab1 and Stab2. Previous transcriptomic analysis of LSECs revealed the most drastic changes in Stab-DKO compared to WT, Stab1-KO and Stab2-KO as well (Olsavszky et al., 2021). Nevertheless, side effects of anti-Stabilin targeted therapies in individuals with decreased renal or liver function cannot be ruled out and should be monitored closely.

To conclude, we were able to further validate our previous findings that the four fasciclin proteins Stab1, Stab2, POSTN and TGFBI constitute a ligand-scavenger receptor system without any relevant compensation by other scavenger receptors. While targeted inhibition of Stab1 or Stab2 might be beneficial in the context of atherosclerosis prevention, double deficiency leads to premature organ failure likely due to multi-ligand depositions caused by deficient hepatic scavenging.

## Supporting information

Supplementary Table 1 containing Antibodies, Probes, Primer

## Acknowledgements

We thank Hiltrud Schönhaber, Maria Muciek, Camela Jost, Stefanie Riester and Janina Ritz for excellent technical support.

