## Supplementary figures and images for "Deficiency for scavenger receptors Stabilin-1 and Stabilin-2 leads to age-dependent renal and hepatic depositions of fasciclin domain proteins TGFBI and Periostin in mice"

### Supp. Figures

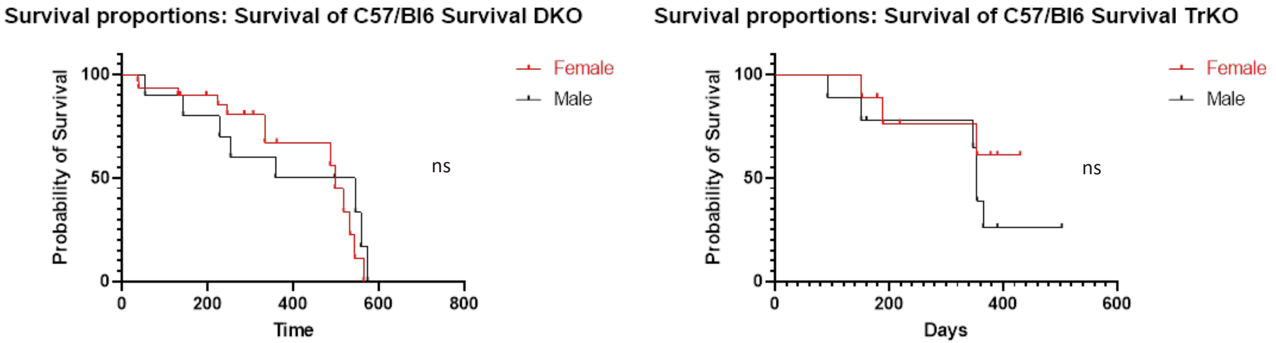

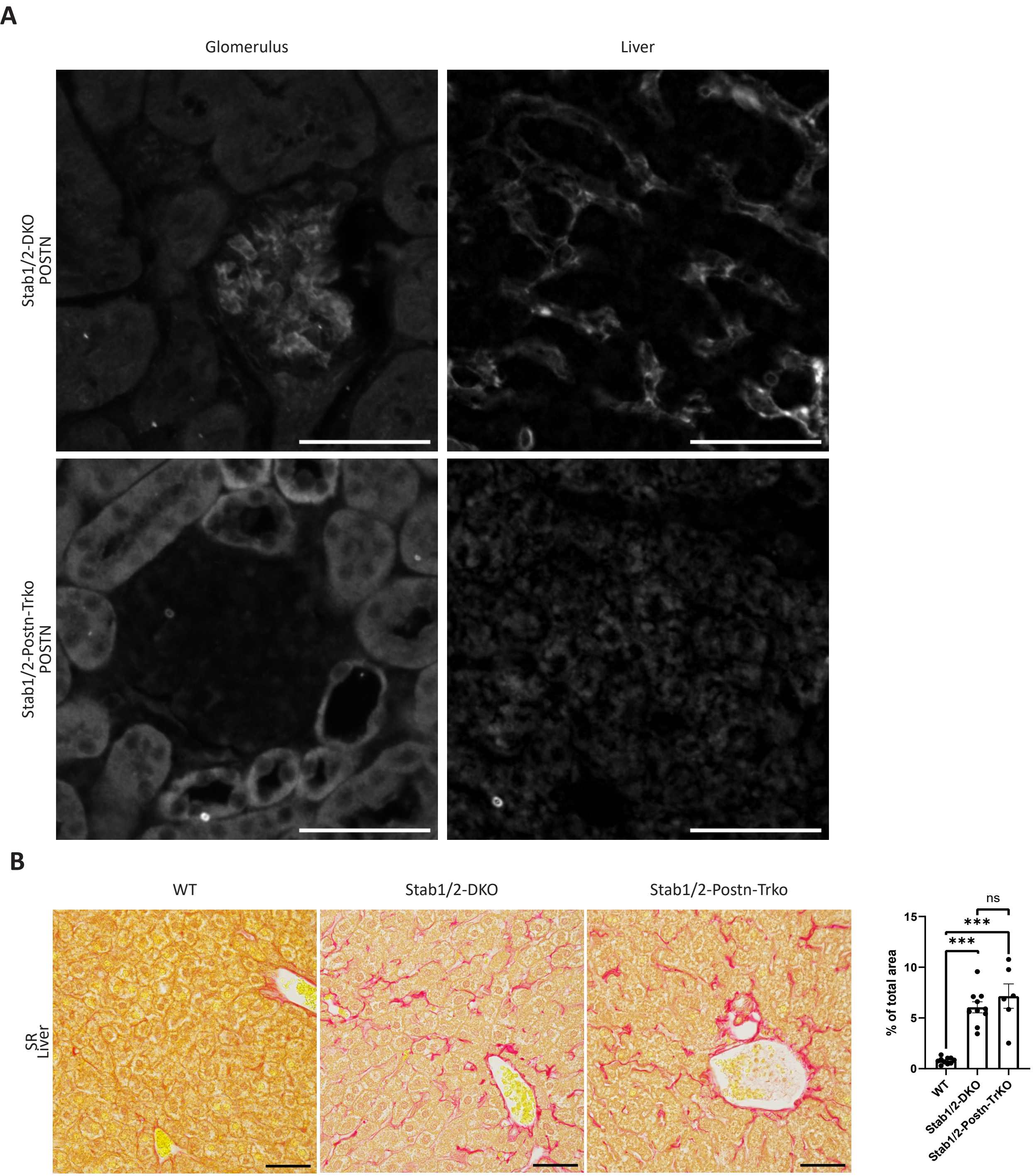

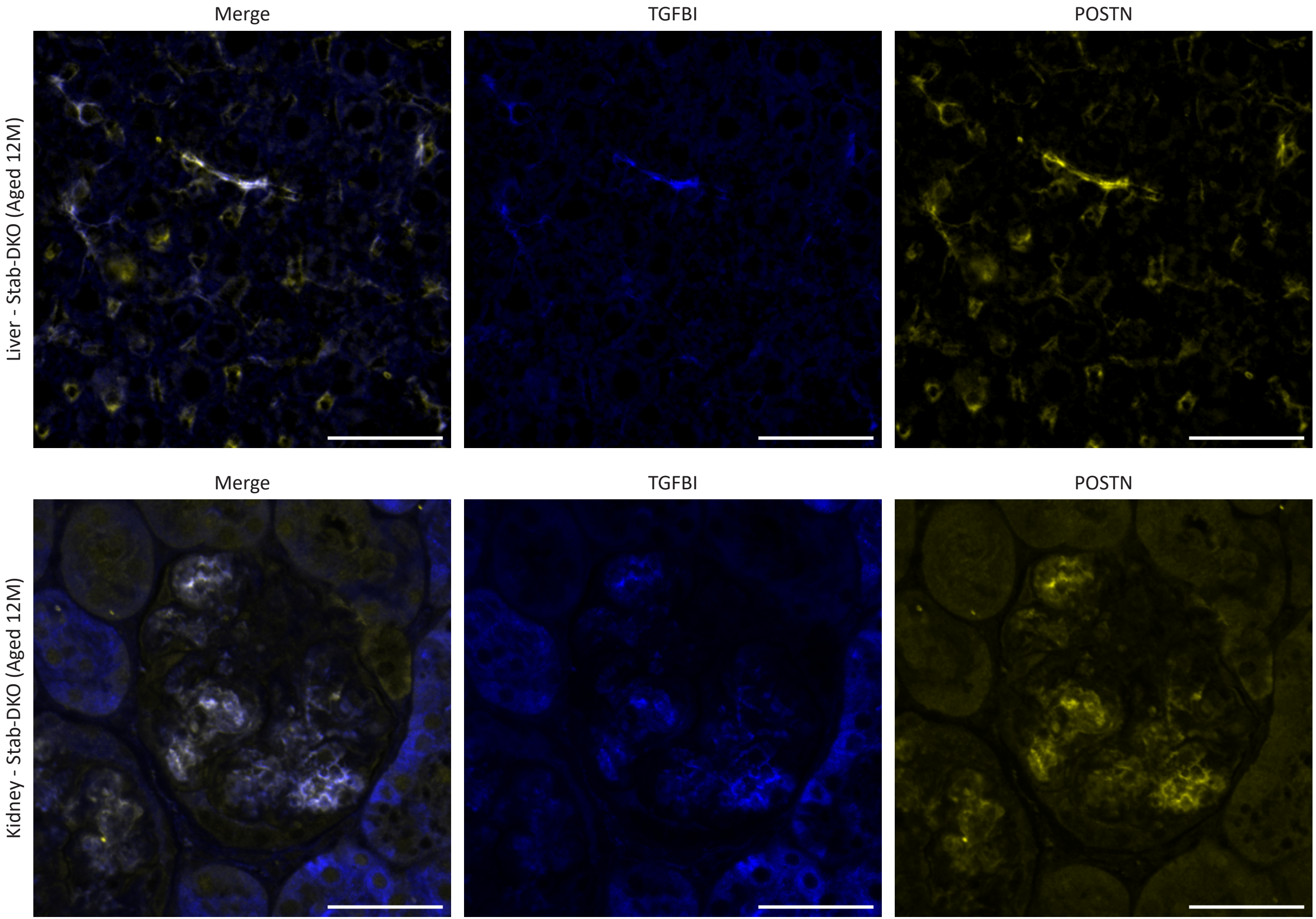

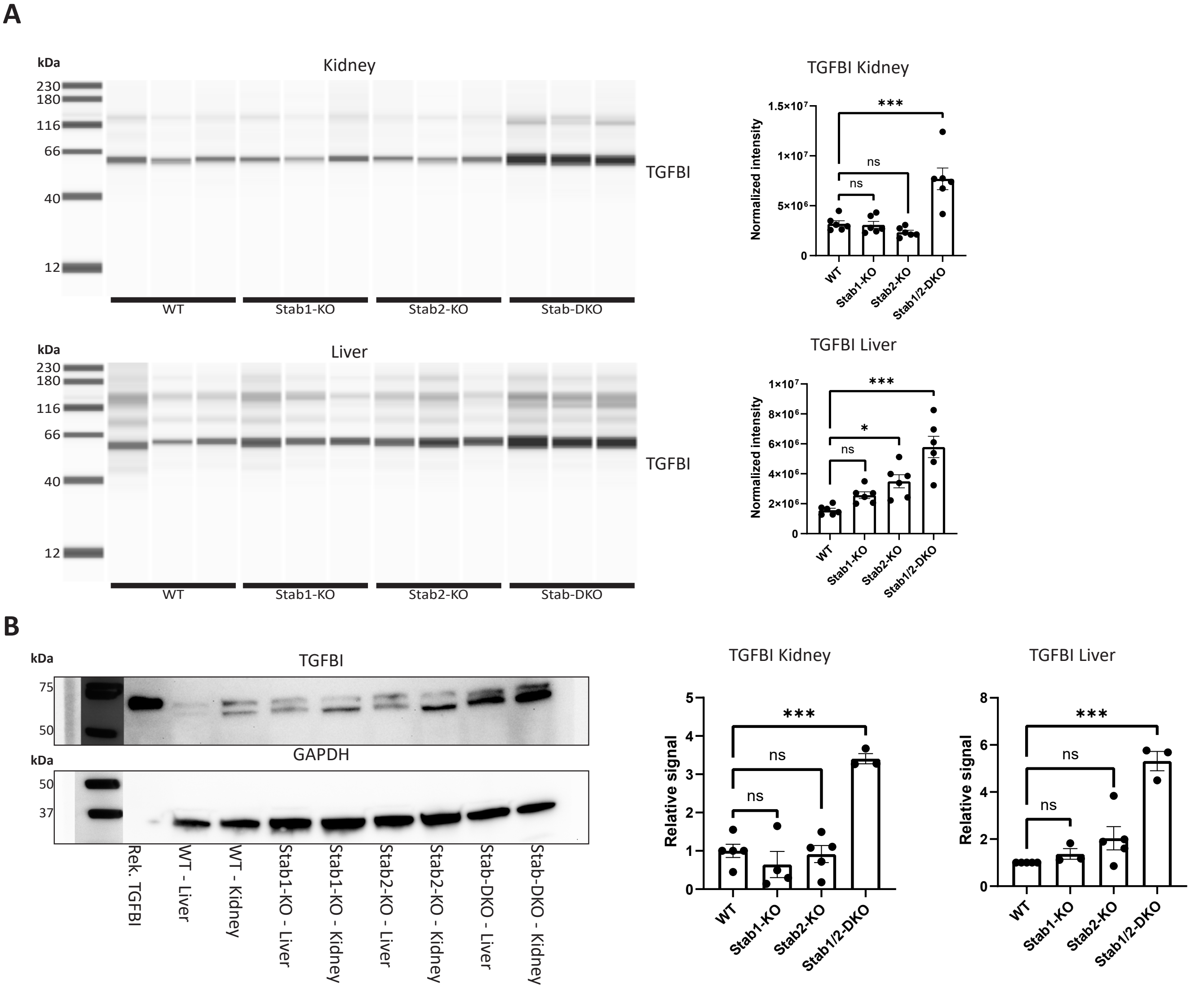
