## Supplementary material for "Deficiency for scavenger receptors Stabilin-1 and Stabilin-2 leads to age-dependent renal and hepatic depositions of fasciclin domain proteins TGFBI and Periostin in mice": Supp. Figure Legends

**S. Fig. 1** Survival curves are similar for both male and female mice

**(A)** Survival curve of male and female Stab-DKO mice (left) and Stab-POSTN-Triple deficient mice (right). n≥5 for all experiments. ns = not significant, *= p<0.05; **=p<0.01; ***=p<0.001.

**S. Fig. 2** Specificity of POSTN antibody in kidney and liver

**(A)** Representative photomicrographs of kidney tissue (left panel) and liver tissue (right panel) of Stab-DKO (upper panel) and Stab-POSTN-Triple deficient mice (lower panel) stained with POSTN. **(B)** Representative photomicrographs of Sirius-red stained liver tissue. Quantification of average Sirius-red positive area of total photomicrograph area is shown on the right (in % of total photomicrograph area). n≥5 for all experiments. Scale bar = 50 µm. ns = not significant, *= p<0.05; **=p<0.01; ***=p<0.001.

**S. Fig. 3** Colocalization of POSTN and TGFBI in liver and kidney tissue

Representative photomicrographs of liver tissue (upper panel) and kidney tissue (lower panel) of Stab-DKO animals stained with TGFBI (blue) and POSTN (yellow). Scale bar = 50 µm

**S. Fig. 4** Simple Western™ and Western blots of kidney and liver tissue from Stabilin-deficient animals

**(A)** Simple Western™ based quantification of TGFBI intensity relative to total protein content in homogenized liver (upper panel) and kidney (lower panel) tissue. N=6 for all experiments. **(B)** Representative photomicrographs of Western-Blot analysis of TGFBI (upper panel) in homogenized kindey and liver tissue. GAPDH was used as a loading control (lower panel), recombinant TGFBI as a positive control (left lane). Quantification of TGFBI intensity relative to GAPDH intensity in homogenized kidney tissue (middle panel) and homogenized liver tissue (right panel) relative to WT. n≥3 for all experiments. ns = not significant, *= p<0.05; **=p<0.01; ***=p<0.001.
